# Rate-Dependent Mechanical Behavior of Human Femoropopliteal Arteries in Biaxial Testing

**DOI:** 10.64898/2026.04.09.717509

**Authors:** Bahman Kargarbahrkhazar, Sayed Ahmadreza Razian, Majid Jadidi

**Author notes:** Correspondence and Reprints requests to: Majid Jadidi, PhD, Department of Biomechanics, University of Nebraska at Omaha.

## Abstract

**Introduction:** Arteries, like other soft tissues, exhibit viscoelastic mechanical behavior, meaning their response to stress and strain is time dependent. This implies that the way arteries deform depends not only on the amount of force applied but also on the rate at which the force is applied. This study investigates the effects of different loading rates on the mechanical behavior of human femoropopliteal arteries (FPAs) to understand their rate-dependent characteristics.

**Methods:** Human FPA specimens were collected from 14 donors, including 7 males and 7 females, aged 45-55 years. A 10x10 mm segment was isolated, mounted onto a biaxial testing device, and subjected to varying loading rates (10 to 50 mN/s). Mechanical responses were recorded, and stress-stretch curves were analyzed. Statistical analyses, including mixed-design ANOVA, assessed the impact of sex and loading rates on tissue stiffness.

**Results:** Results indicated significant loading-rate dependency, particularly in the circumferential direction. Stretch values decreased with increasing loading rates, more prominently in the circumferential than in the longitudinal direction (p-value<0.01). Statistical analyses revealed no significant interaction between sex and loading rate, though male arteries exhibited slightly higher compliance than female arteries.

**Discussion:** The findings demonstrate that the mechanical response of FPAs is highly dependent on the loading rate, with more pronounced effects observed in the circumferential direction. At higher loading rates, the human FPAs demonstrated a stiffer response in the circumferential direction.

**Dedication:** We dedicate this work to the memory of our late student, Ali Zolfaghari Sichani, who passed away tragically during his doctoral studies. Ali performed the majority of the experiments and the initial analysis reported in this paper. His passion, dedication, and hard work were the foundation of this research, and he is deeply missed.

## 1 Introduction

Arteries, similar to other soft tissues, exhibit viscoelastic mechanical behavior, meaning their response to stress and strain is time-dependent^1–4^. This viscoelasticity implies that the way arteries deform depends not only on the amount of force applied but also on the rate at which the force is applied. One crucial aspect of viscoelasticity is loading-rate dependency, which significantly affects arterial behavior^5,6^. Despite the inherent viscoelastic nature of arteries, they are often modeled using the pseudoelastic assumption for simplicity^2,7–9^. Pseudoelasticity approximates the mechanical response of viscoelastic materials by treating them as elastic during analysis, thereby simplifying the complexity of the model. This simplification, while not capturing all aspects of arterial behavior, provides useful insights into their mechanical properties under different loading conditions.

To assess the mechanical properties of arteries, researchers have adopted various methods, with planar biaxial extension testing being widely used^10–12^. In this approach, arterial samples are opened along their length and mounted onto a biaxial testing device, aligning the circumferential and longitudinal directions of the tissue with the testing axes. The sample is then subjected to a series of stretches or forces to evaluate its mechanical properties. In stretch-controlled tests, a maximum stretch is defined for both directions, and the tissue is stretched to these limits at a specified strain rate. Preconditioning cycles, typically ranging from 5 to 20, are often applied to stabilize the mechanical response of the tissue before the main testing and assume pseudoelastic response.

Previous studies have utilized different strain rates in biaxial testing of soft tissues. For instance, Jadidi et al. used a stretch rate of 1%s^−1^ to determine the mechanical properties of aorta, iliac, and femoropopliteal arteries^13–16^. Sommer et al. employed a stretch rate of 3 mm/min to study the left ventricles of human hearts^17^. Vélez-Rendón et al. performed planar biaxial mechanical testing on the right ventricles of rats with 10% stretch-controlled tests at 0.5 Hz^18^. Terry et al. tested biaxial tissue samples from swine at a mean rate of 0.2 Nm/s using load-controlled experiments^19^.

Previous research has shown varied responses in soft tissues under different loading rates, with some studies reporting stiffening and others observing softening or no significant change in mechanical properties. These discrepancies may arise from differences in loading rates, tissue types, and test conditions. Understanding the specific mechanical behavior of human arteries under different loading rates is crucial, as these insights can guide computational studies using the material parameters derived from the mechanical tests.

Our study focuses on the load-rate dependency of the human FPA. As a muscular artery, the FPA has a distinct structure compared to the aorta and other blood vessels^20^, suggesting its rate dependency might differ. By examining the mechanical behavior of human FPAs under various loading rates, this research aims to fill gaps in our understanding of FPA biomechanics.

## 2 Methods

### 2.1 Specimens

Human FPA specimens were obtained from 14 donors aged 45-55 years (7 males, 7 females, average age 50±3 years) to minimize age-related variability in arterial properties (Table 1) ^21^. The specimens were procured by the organ procurement organization, Live On Nebraska, within 24 hours of death, with next of kin consent. The tissues were transported to our lab at 4°C in 0.9% phosphate-buffered saline (PBS) to preserve freshness. Arteries were then dissected and cleaned of surrounding tissues.

**Table 1.**
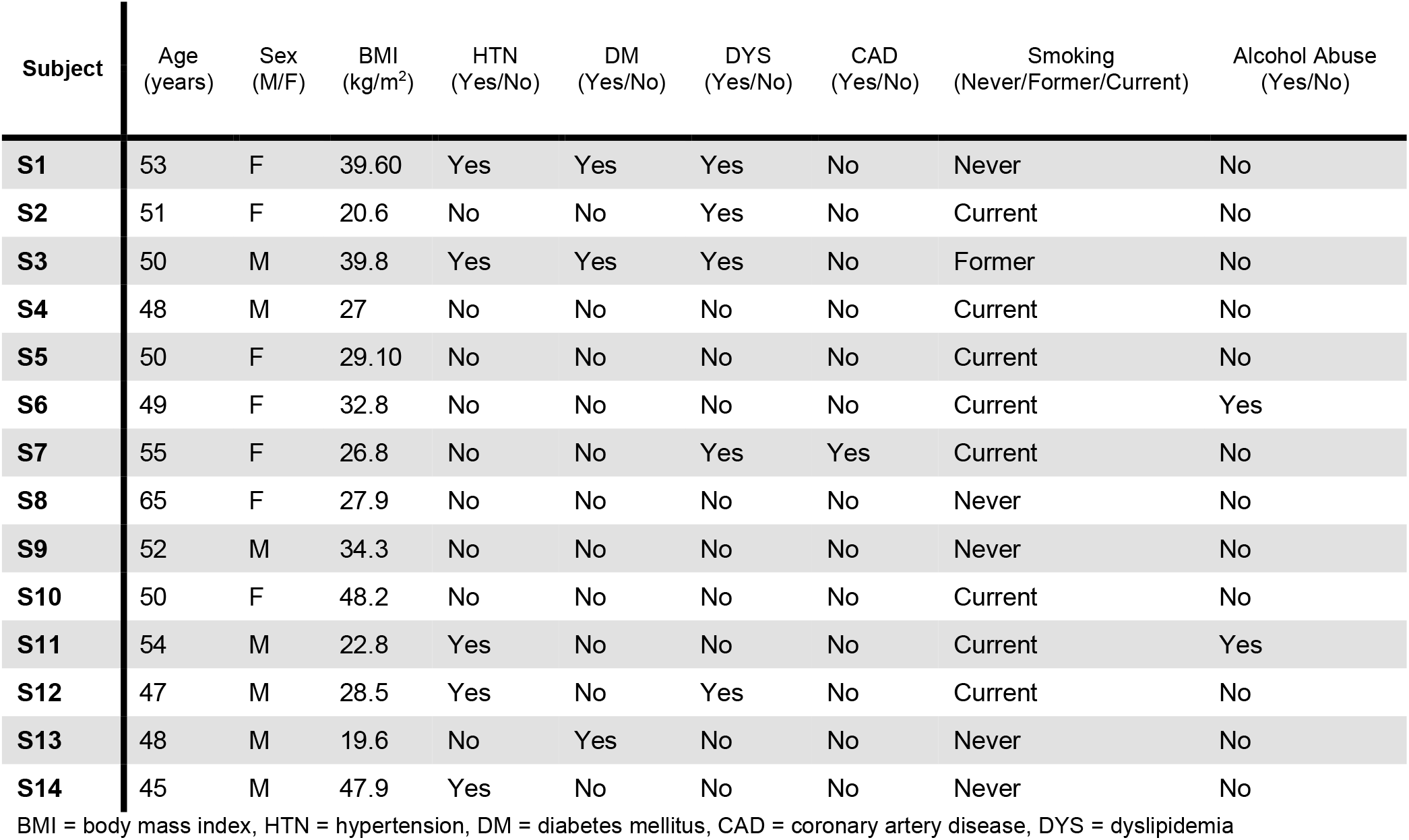
Subject Demographics and risk factors.

A 13 mm segment of the artery was isolated from the proximal FPA, right below the profunda femoris artery bifurcation. This segment was cut longitudinally and a 10x10 mm square specimen was separated for the biaxial test. Post-test completion, three axial strips were obtained from the middle of the specimen, and photographed using a Nikon z7 camera to measure thickness of the specimen for further stress calculations. We determined the wall thickness by selecting a series of equally spaced points on the intimal and adventitial surface of each of the strips and calculating the distance between these points, providing the average wall thickness for each axial strip. The average thickness for each specimen was then determined by averaging the thickness of the three axial strips.

### 2.2 Planar Biaxial Test

Planar biaxial tests were conducted using a CellScale biaxial device equipped with 2.5 N load cells. The 10x10 mm square specimen was attached such that the circumferential direction of the artery aligned with the biaxial device’s vertical axis (index *θ*), and the artery’s longitudinal direction aligned with the biaxial device’s horizontal axis (index *z*). The samples were immersed in a PBS bath maintained at approximately 37°C during testing (Figure 1).

**Figure 1:**
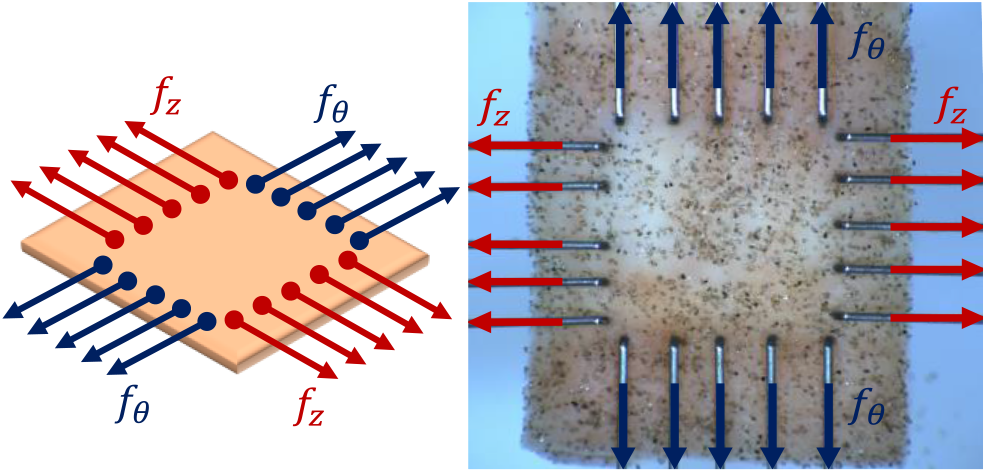
Left: Schematic representation of the planar biaxial test. Right: Experimental setup showing a sample undergoing stretching with applied forces. *f*_θ_and *f*_*z*_ are the forces in the circumferential and longitudinal directions of the arterial wall specimen, respectively.

Following the methodology outlined in a previous study^22^, which demonstrated the negligible effect of shear for FPAs, the samples were attached using BioRake. A sparse distribution of sand particles was applied to the samples, and images and data were recorded and saved at a rate of 5 Hz. Upon completing the tests, these images were processed for displacement analysis. The sand particles served as visual markers, allowing for the tracking of movement and measurement of sample displacement during stretching.

The protocols for the load-controlled tests at specific load rates are detailed in Table 2. Prior to initiating these protocols, the samples underwent 10 preconditioning cycles at a rate of 10 mN/s to establish a consistent stress-stretch response. Subsequently, the load-controlled protocol was executed in sequence from the slowest to the highest rate, i.e., 10 to 50 mN/s, with a maximum force of 800 mN applied in both directions. The selection of this maximum force was based on previous studies which indicated that this level of force induces non-linearity in the force-stretch curves while preventing excessive stretching that could potentially lead to tissue damage^23^.

**Table 2.** Load-controlled test protocols for each load rate after preconditioning cycles.

| Protocol | Ratio | Maximum Force Magnitude (mN) |  |
| --- | --- | --- | --- |
|  |  | Longitudinal | Circumferential |
| Equi-Biaxial | 1.0:1.0 | 800 | 800 |
| Ratio-X | 0.5:1.0 | 400 | 800 |
| Equi-Biaxial | 1.0:1.0 | 800 | 800 |
| Ratio-Y | 1.0:0.5 | 800 | 400 |
| Equi-Biaxial | 1.0:1.0 | 800 | 800 |

Following the initial preconditioning phase, the protocols outlined in Table 2 were repeated at five different loading rates: 10, 20, 30, 40, and 50 mN/s. As shown in this table, three distinct protocols were employed. In the Equi-Biaxial protocol, the sample undergoes equal stretching in both directions, reaching a maximum load of 800 mN. Under the Ratio-X protocol, the sample is stretched to 400 mN longitudinally and 800 mN circumferentially. Conversely, in the Ratio-Y protocol, the sample is stretched 400 mN circumferentially and 800 mN longitudinally.

### 2.3 Stress-Stretch Curve

Image processing was employed to measure the deformation gradient at the center of the specimen, away from the rakes. Initially, four points were identified on the specimen in the first image. Using a data analysis software integrated within the CellScale software package, these four points were tracked across all images recorded by the biaxial device’s camera. The coordinate positions of these points were then exported into a file, providing their coordinates over time for all protocols. Since data and images were collected at 5 Hz, there are 5 datapoints in each second. Furthermore, another file was exported containing force values recorded in both directions by the load cells. Using the tracking data, we determined the stretches experienced by the specimen. Subsequently, these stretches and the forces measured by the device were used to compute the Cauchy stresses in the circumferential (*T*_*θθ*_) and longitudinal (*T*_*zz*_) directions. The stresses were calculated as follows:

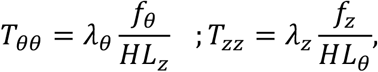

In these equations *f*_*θ*_ and *f*_*z*_ are the forces in the circumferential and longitudinal direction, respectively, *H* represents the stress-free thickness of the specimen, and *L*_*θ*_ and *L*_*z*_ are the initial dimensions of the specimen in the circumferential and longitudinal directions, respectively^12,24^. In calculating the stress-stretch data, a reference point is essential for determining displacements and stretches. One common reference is the constant reference point, where the first image after the end of preconditioning protocol is selected as the reference, and other stretches are calculated relative to this image. In this case, since we performed a load-controlled test, the specimens will have some residual deformation at zero load at the end of each protocol. Therefore, the stretch at the beginning of each protocol starts from a value greater than 1. Another possible reference configuration is the variable reference, where stretches for each protocol are calculated from the beginning of the protocol. In this case, the stretch value of each protocol starts from 1.0. The difference between the constant and variable reference configuration is illustrated in Figure 2. In our study, we utilized the variable reference configuration.

**Figure 2:**
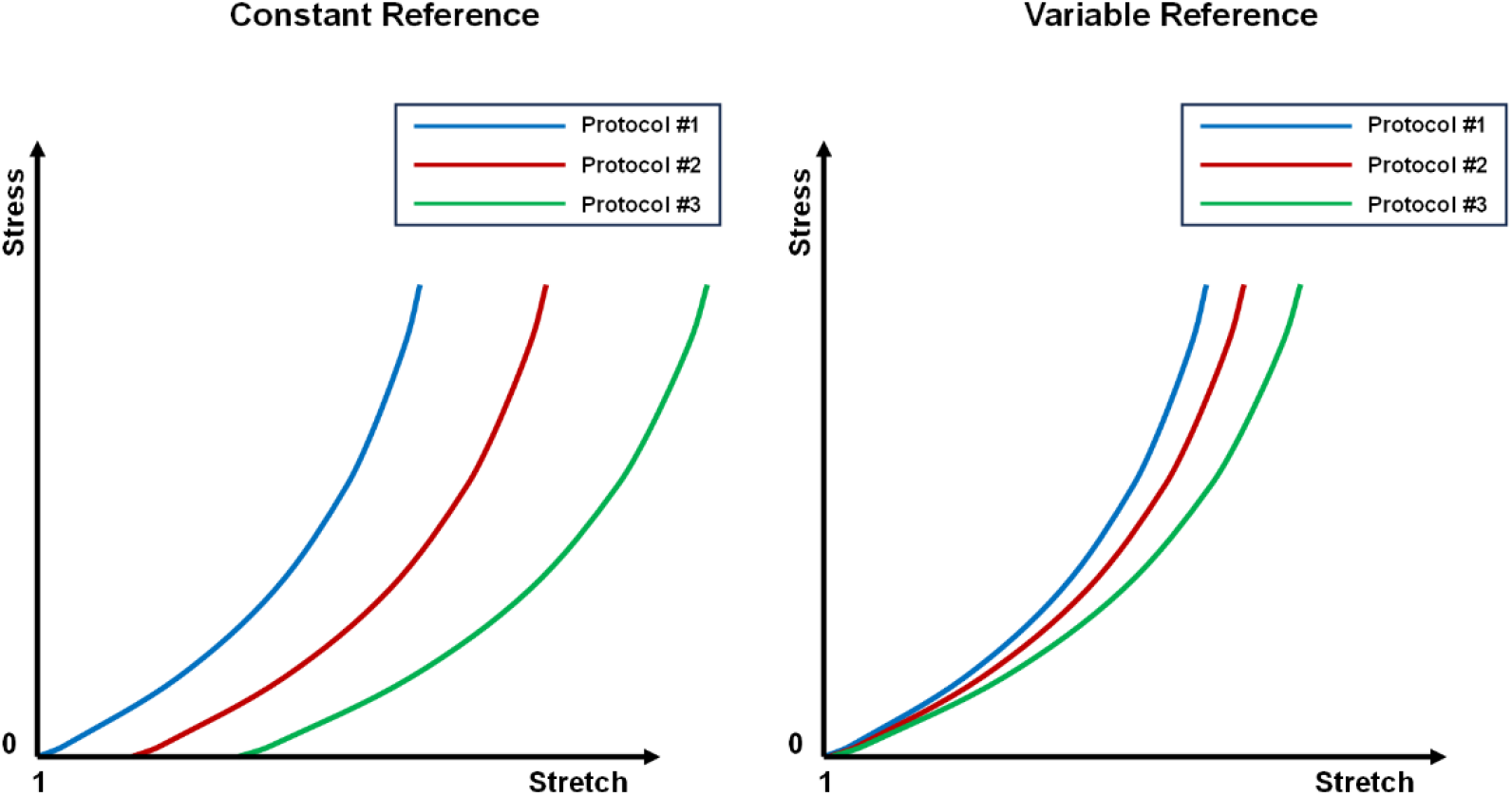
Illustration of two reference configurations for calculating stress and stretch values: the constant reference configuration (left) and the variable reference configuration (right).

### 2.4 Statistical Analysis

To compare the samples statistically, the stretches corresponding to the maximum stress level were determined for all samples. In the Equi-Biaxial protocol, the maximum stress level was consistently 70 kPa in both directions for all samples. Stretch values for each sample under five different speed rates were calculated at specific stress levels: 70, 50, and 30 kPa. Additionally, we extracted and compared results from three tested protocols: Equi-Biaxial, Ratio-X, and Ratio-Y.

To assess the significance of the observed differences, we did different statistical tests. Considering that the data comprises repeated measurements involving two factors (sex as the between-subject factor and different load rates as the within-subject factor), we opted for a mixed design ANOVA test. Due to the repeated measurements, we also considered the assumption of sphericity, which requires the variances of the differences between all combinations of measurement groups to be equal. Degrees of freedom were adjusted using the Greenhouse-Geisser correction method. If a significant p-value (p < 0.05) is observed in the ANOVA test, it indicates significant differences between the group means. In addition to the tests of within-subject effects, we conducted pairwise comparisons between different groups using the Bonferroni method to determine which specific groups were significantly different.

## 3 Results

Figure 3 provides a visual representation of the mechanical behavior of a representative FPA sample (S3, 50-year-old male) under varying loading rates in two directions. The stretch values in the longitudinal direction consistently exceed those in the circumferential direction across all loading rates. Additionally, the stretch values decrease at different stress levels as the loading rate increases as the stress-stretch curves shift toward left. This pattern is observed in both the longitudinal and circumferential directions. The difference in stretch values is particularly noticeable when comparing the rates of 10 mN/s and 50 mN/s. For this sample, under the Equi-Biaxial protocol, the stretch values decreased from 1.26 to 1.23 in the longitudinal direction and from 1.12 to 1.10 in the circumferential direction at a stress level of 70 kPa. Supplement file provides all the stress-stretch curves of the Equi-Biaxial protocol for all samples in both directions.

**Figure 3:**
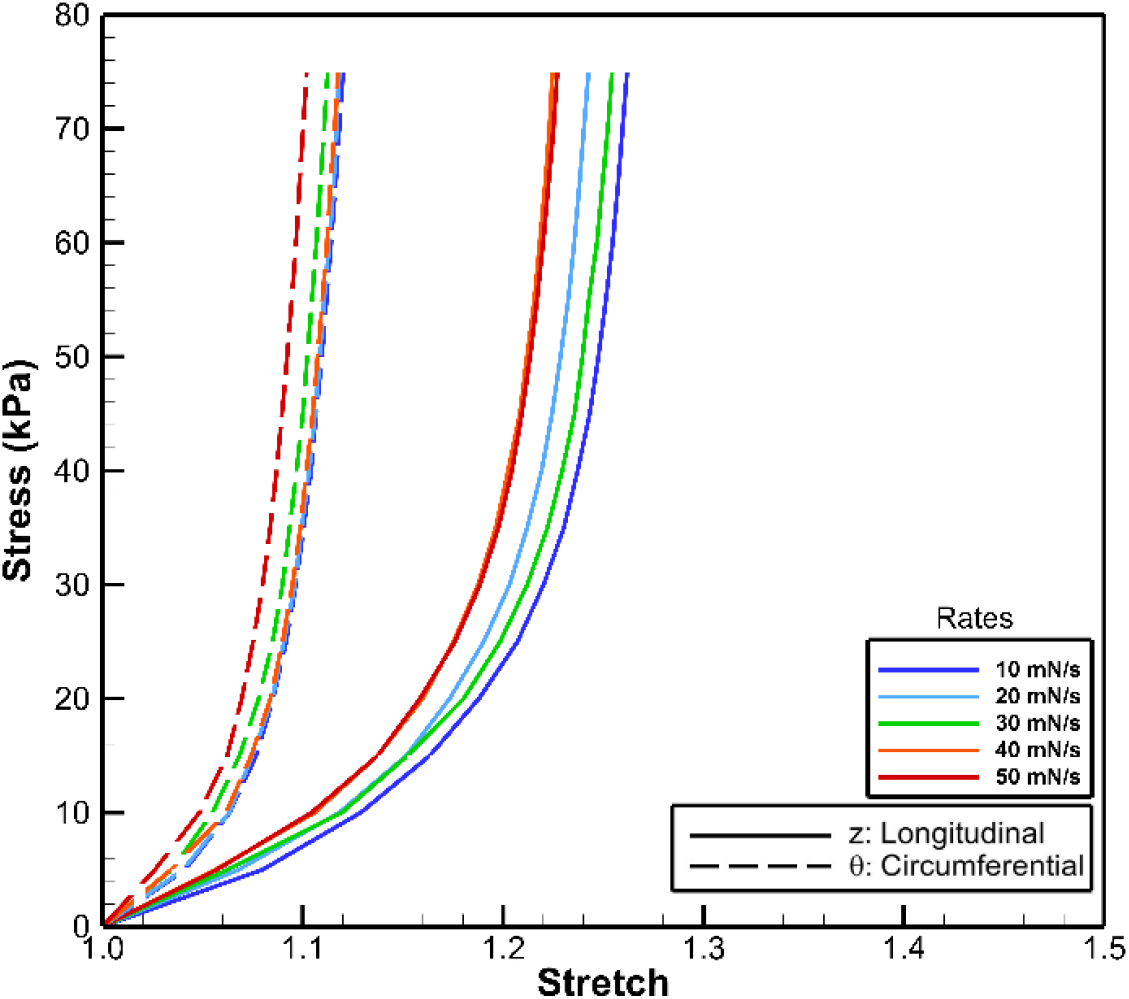
Equi-Biaxial stress-stretch curves of S3 (50-year-old male) sample in both the longitudinal and circumferential directions at different loading rates.

We have three protocols and three stress levels in both longitudinal and circumferential directions for ANOVA analysis. Each protocol is investigated separately to determine the effect of the loading rates on the stretch values. Figure 4 shows the mean and standard deviation of the stretch values for the Equi-Biaxial protocol at specific stress levels. The red lines represent the female samples, and the blue lines represent the male samples. Additionally, these figures display the results for both longitudinal (solid lines) and circumferential (dashed lines) directions. As observed, the mean stretch values for males are consistently higher than those for females across all stress levels. Additionally, the stretch values in the circumferential direction are lower than those in the longitudinal direction for both males and females across all stress levels. Male arteries appear to be slightly more compliant, as they experience higher stretches at the same level of stress.

**Figure 4:**
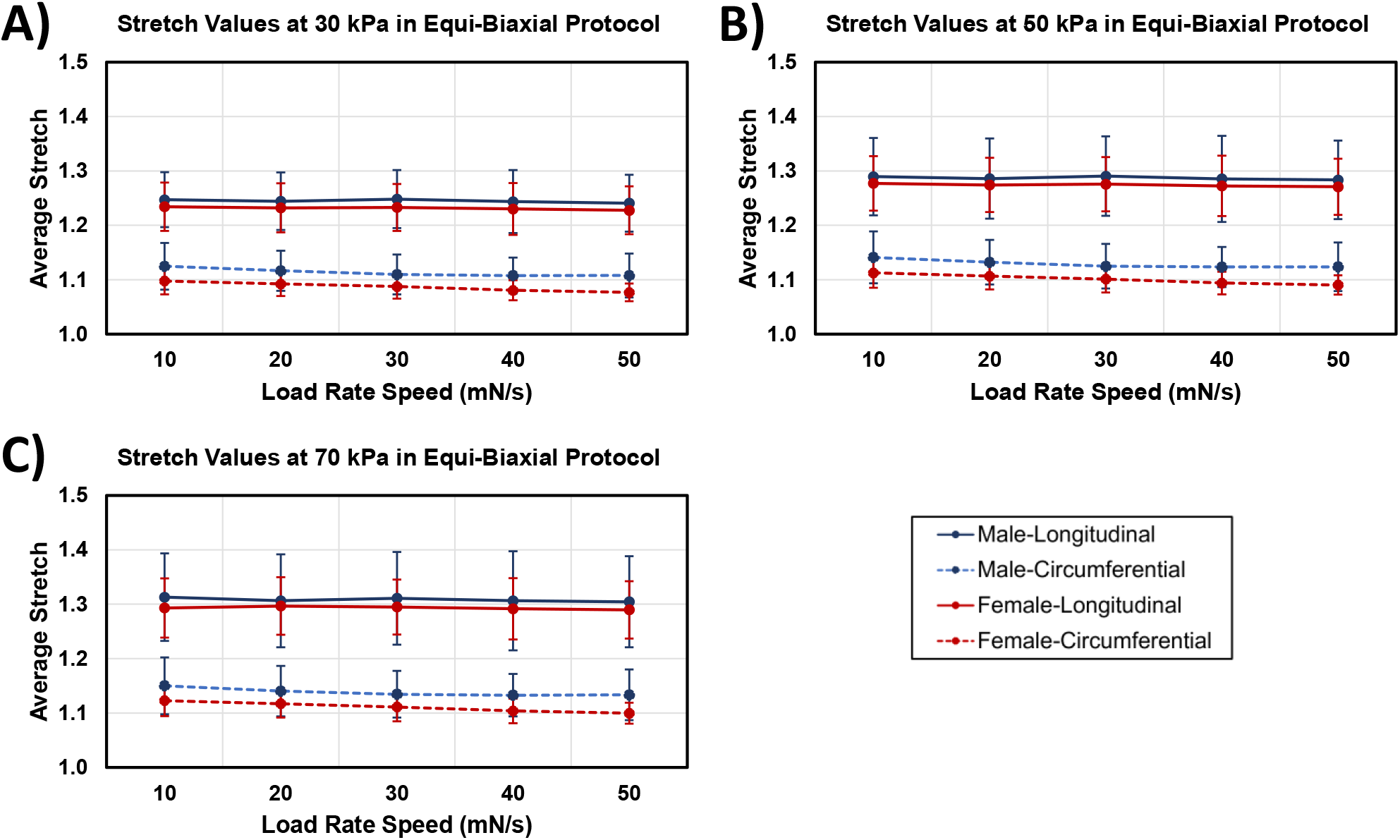
Descriptive plots showing the stretch values of the Equi-Biaxial protocol at three different stress levels for both sexes in two directions across five different loading rates. A) Stretch values at the stress level of 30 kPa. B) Stretch values at the stress level of 50 kPa, and C) Stretch values at the stress level of 70 kPa.

At 30 kPa in the Equi-Biaxial protocol, the mean stretch values for all rates in the longitudinal direction are 1.24 ± 0.05 for males and 1.23 ± 0.04 for females. In the circumferential direction, the mean stretch values for males decreased from 1.12 ± 0.04 to 1.10 ± 0.04, and for females, it decreased from 1.10 ± 0.03 to 1.08 ± 0.02 with increasing loading rates.

At 50 kPa in the Equi-Biaxial protocol, the mean stretch values for all rates in the longitudinal direction are 1.28 ± 0.07 for males and 1.27 ± 0.05 for females. In the circumferential direction, the mean stretch values for males decreased from 1.14 ± 0.04 to 1.12 ± 0.04, and for females, it decreased from 1.11 ± 0.03 to 1.09 ± 0.02 with increasing loading rates.

At 70 kPa in the Equi-Biaxial protocol, the mean stretch values for all rates in the longitudinal direction are 1.31 ± 0.09 for males and 1.30 ± 0.05 for females. In the circumferential direction, the mean stretch values for males decreased from 1.15 ± 0.05 to 1.13 ± 0.05, and for females, it decreased from 1.12 ± 0.03 to 1.10 ± 0.02 with increasing loading rates.

To compare the results of the Equi-Biaxial protocol, we used ANOVA tests. We computed a mixed-design ANOVA to determine the effects of sex and load rate on tissue stiffness. Figure 4 provides the means and standard deviations for each condition. Additionally, all data were checked for normality and homogeneity, and our data met these assumptions, allowing us to use parametric tests. Sphericity was also checked, and because this assumption was violated, the results were adjusted using the Green-Geisser method.

Table 3 shows the p-values and other important statistical analyses. The results of stretches in the Equi-Biaxial protocol indicate that sex and the interaction between sex and load rate are not significant in either direction or at any of the three stress levels due to the low F ratios and high p-values. Therefore, we focused on the main effect of load rate. In the longitudinal direction, the F ratio for all stress levels is below 1.0, and the p-values are considerably higher than 0.3, indicating insufficient evidence to conclude that the load rate affects the stretch values at different speeds. However, in the circumferential direction, the load rate effect has a high F ratio and significant p-values at all three stress levels, indicating that the load rate affects the stretch values in the circumferential direction.

**Table 3.**
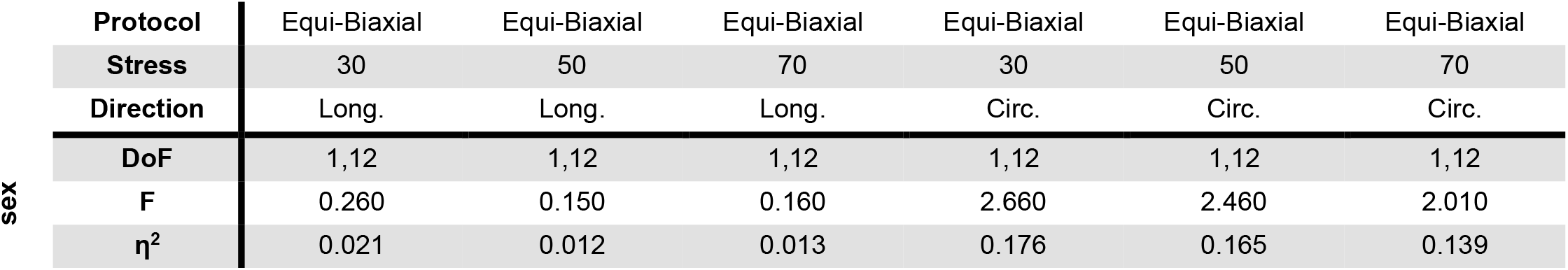
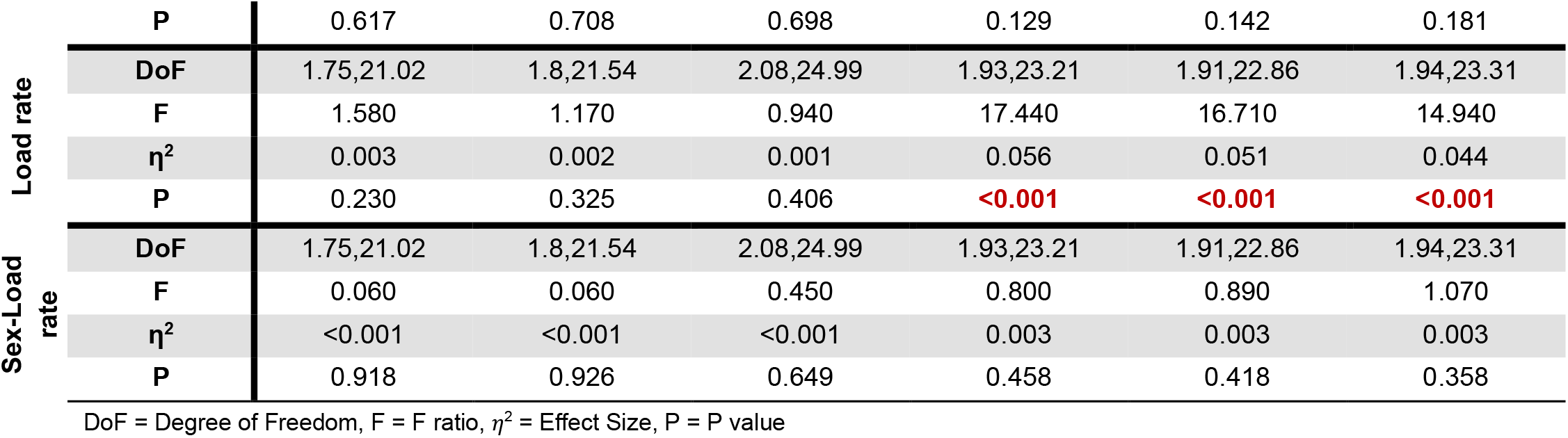
ANOVA table for the Equi-Biaxial protocol in different directions.

In Figure 5, the mean stretch values of the Ratio-X protocol are shown. Since, in the Ratio-X protocol, the specimens are only stretched to a 400 mN load in the longitudinal direction, the maximum stress level is lower than that in the Equi-Biaxial protocol. Therefore, we compared the stretch values at 30 kPa. At this stress level, the mean stretches in the longitudinal direction exceed those in the circumferential direction. Furthermore, the stretch values for male subjects are higher than those for female subjects in the Ratio-X protocol.

**Figure 5:**
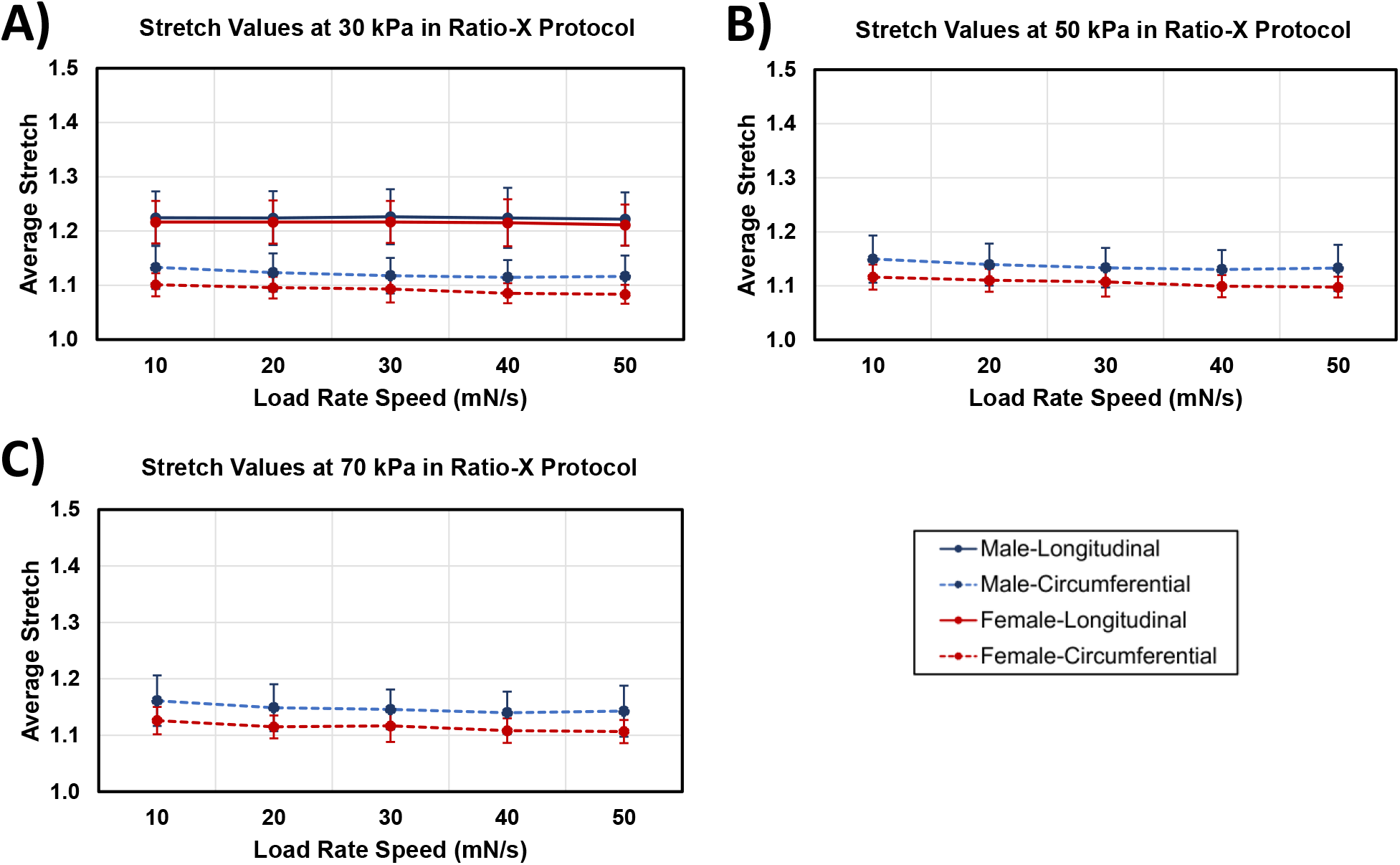
Descriptive plots showing the stretch values of the Ratio-X protocol at three different stress levels for both sexes in two directions across five different loading rates. A) Stretch values at the stress level of 30 kPa in both directions. B) Stretch values at the stress level of 50 kPa in the circumferential direction, and C) Stretch values at the stress level of 70 kPa in the circumferential direction.

At 30 kPa in the Ratio-X protocol, the mean stretch values in the longitudinal direction for all rates are 1.22 ± 0.05 for males and 1.21 ± 0.04 for females. In the circumferential direction, the mean stretch values for males decreased from 1.13 ± 0.04 to 1.11 ± 0.04, and for females, it decreased from 1.10 ± 0.02 to 1.08 ± 0.02 with increasing loading rates. At 50 kPa in the circumferential direction, the mean stretch values for males decreased from 1.15 ± 0.04 to 1.13 ± 0.04, and for females, it decreased from 1.12 ± 0.03 to 1.10 ± 0.02 with increasing loading rates. At 70 kPa in the circumferential direction, the mean stretch values for males decreased from 1.16 ± 0.04 to 1.14 ± 0.04, and for females, it decreased from 1.13 ± 0.02 to 1.11 ± 0.02 with increasing loading rates.

In Figure 6, the results of the mean stretch values for the Ratio-Y protocol are displayed. In the Ratio-Y protocol, the maximum applied load is 400 mN in the circumferential direction, resulting in a maximum stress level lower than that of the Equi-Biaxial protocol. Accordingly, the stretch values at 30 kPa are compared in this direction. Similar to the two previous protocols, male subjects exhibit higher mean stretch values in both directions compared to female subjects. Additionally, at the stress level of 30 kPa, the longitudinal stretch values exceed those of the circumferential direction for each sex and load rate.

**Figure 6:**
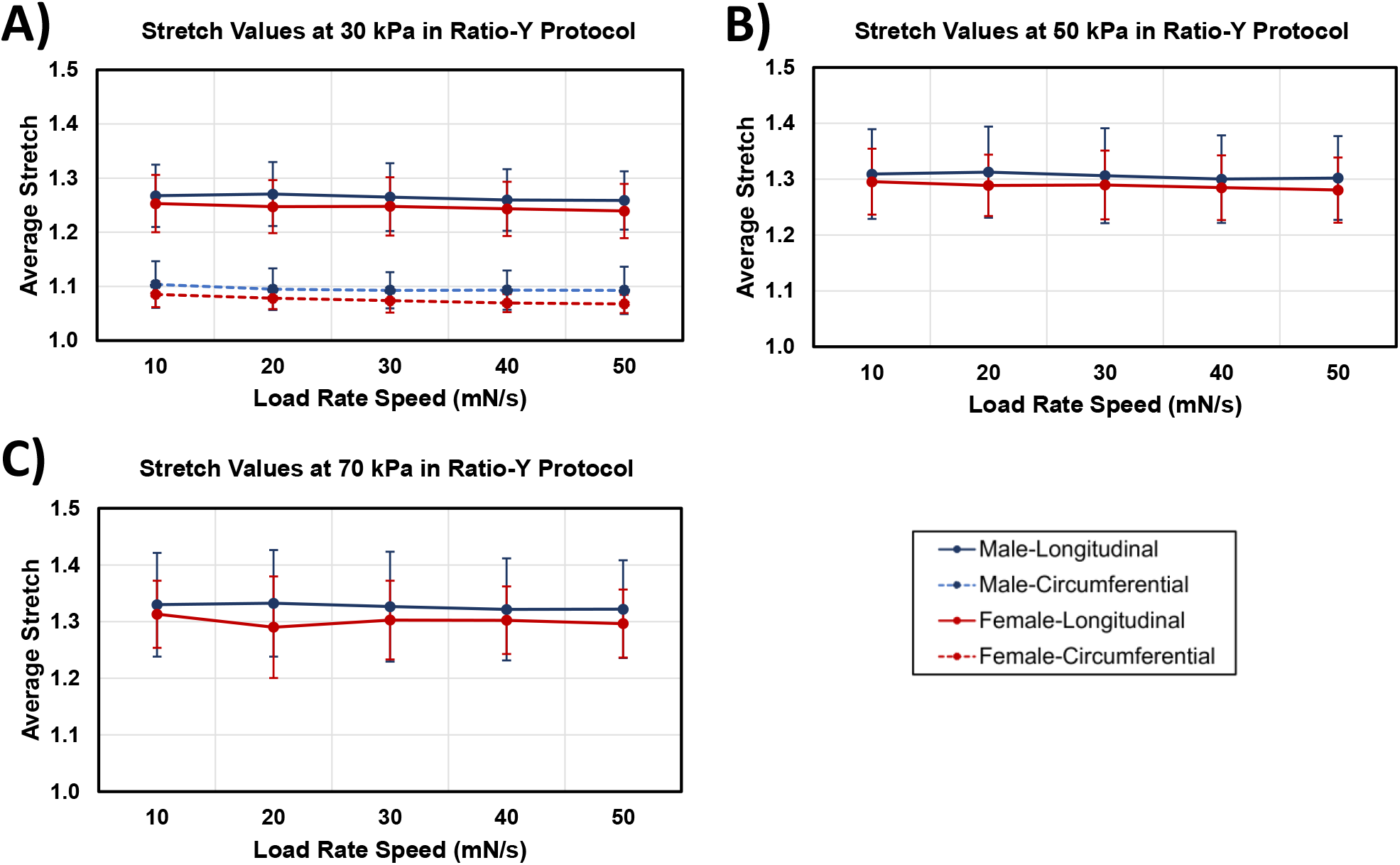
Descriptive plots showing the stretch values of the Ratio-Y protocol at three different stress levels for both sexes in two directions across five different loading rates. A) Stretch values at the stress level of 30 kPa in both directions. B) Stretch values at the stress level of 50 kPa in the longitudinal direction, and C) Stretch values at the stress level of 70 kPa in the longitudinal direction.

At 30 kPa in the Ratio-Y protocol, the mean stretch values in the longitudinal direction for all rates are 1.26 ± 0.05 for males and 1.25 ± 0.05 for females. In the circumferential direction, the mean stretch values for males decreased from 1.10 ± 0.04 to 1.09 ± 0.04, and for females, it decreased from 1.09 ± 0.02 to 1.07 ± 0.02 with increasing loading rates. At 50 kPa in the longitudinal direction, the mean stretch values for all rates are 1.30 ± 0.08 for males and 1.29 ± 0.06 for females. At 70 kPa in the longitudinal direction, the mean stretch values for all rates are 1.33 ± 0.09 for males and 1.31 ± 0.07 for females.

In Figure 5 and Figure *6*, the means and standard deviations for the Ratio-X and Ratio-Y protocols are shown. To compare these results, a mixed-design ANOVA test was performed. All the data met the assumptions of normality and homogeneity. The ANOVA results were adjusted using the Green-Geisser method. Table 4 presents the p-values and other important statistical analyses.

**Table 4.** ANOVA table for the Ratio protocols in different directions.

|  | Protocol | Ratio-X | Ratio-X | Ratio-X | Ratio-X | Ratio-Y | Ratio-Y | Ratio-Y | Ratio-Y |
| --- | --- | --- | --- | --- | --- | --- | --- | --- | --- |
|  | Stress | 30 | 30 | 50 | 70 | 30 | 50 | 70 | 30 |
|  | Direction | Long. | Circ. | Circ. | Circ. | Long. | Long. | Long. | Circ. |
| sex | DoF | 1,12 | 1,12 | 1,12 | 1,12 | 1,12 | 1,12 | 1,12 | 1,12 |
|  | F | 0.140 | 3.720 | 3.380 | 3.690 | 0.390 | 0.240 | 0.350 | 1.610 |
| | $\eta^2$ | 0.011 | 0.229 | 0.213 | 0.225 | 0.031 | 0.019 | 0.028 | 0.114 |
|  | P | 0.715 | 0.078 | 0.091 | 0.079 | 0.543 | 0.633 | 0.562 | 0.228 |
| Load rate | DoF | 1.82,21.80 | 2.67,32.08 | 2.70,32.39 | 2.93,35.12 | 1.84,22.06 | 1.96,23.52 | 2.18,26.19 | 2.58,30.91 |
|  | F | 0.930 | 17.340 | 16.870 | 11.680 | 4.480 | 3.630 | 1.280 | 7.760 |
| | $\eta^2$ | 0.002 | 0.054 | 0.048 | 0.049 | 0.007 | 0.005 | 0.003 | 0.030 |
|  | P | 0.401 | <0.001 | <0.001 | <0.001 | 0.026 | 0.043 | 0.296 | <0.001 |
| Sex-Load rate | DoF | 1.82,21.80 | 2.67,32.08 | 2.70,32.39 | 2.93,35.12 | 1.84,22.06 | 1.96,23.52 | 2.18,26.19 | 2.58,30.91 |
|  | F | 0.130 | 0.970 | 1.010 | 0.360 | 0.560 | 0.710 | 1.430 | 0.810 |
| | $\eta^2$ | <0.001 | 0.003 | 0.003 | 0.002 | <0.001 | <0.001 | 0.004 | 0.003 |
|  | P | 0.861 | 0.412 | 0.395 | 0.778 | 0.564 | 0.499 | 0.257 | 0.481 |

DoF = Degree of Freedom, F = F ratio, $\eta^2$ = Effect Size, P = P value

Similar to the Equi-Biaxial protocol, neither sex nor the interaction between sex and load rate show significant effects, indicated by their low F ratios and high p-values in both directions and at all stress levels. Regarding the load rate factor, we observed that in the longitudinal direction, the p-value is not significant. However, in the circumferential direction, the p-value is lower than 0.001. In the Ratio-X protocol, the high F ratios and low p-values provide sufficient evidence that load rates affect the stretch values in this protocol, particularly in the circumferential direction across all three stress levels. Similarly, we observe the same pattern for the Ratio-Y protocol at the stress level of 30 kPa in the circumferential direction.

After conducting pairwise comparisons for the conditions with significant p-values, differences between the load rates were observed. Table 5 illustrates the variations in stretches between each load rate for males and females separately in the circumferential direction. The positive symbol (+) indicates a significant p-value between the specified load rates, while the negative symbol (-) denotes no differences. The observed differences primarily exist in the circumferential direction, and no significant p-values were found for the longitudinal direction, as mentioned in previous sections. All positive comparisons suggest that the stretch values for the higher load rate exceed those for the lower load rate. Notably, for load rate intervals close in value (e.g., 10-20, 20-30, 30-40, 40-50), no differences were observed for both females and males across all mentioned protocols in the table. Additionally, for load rate intervals 20-40, 20-50, and 30-50 in males, no differences were observed.

**Table 5.** Pairwise comparisons between different rates in different protocols. The positive symbol (+) indicates significance between load rates, while the negative symbol (-) indicates no difference.

| Protocol |  | Equi-Biaxial | Equi-Biaxial | Equi-Biaxial | Equi-Biaxial | Ratio-X | Ratio-X | Ratio-X | Ratio-Y |
| --- | --- | --- | --- | --- | --- | --- | --- | --- | --- |
| Stress |  | 30 | 50 | 70 | 30 | 30 | 50 | 70 | 30 |
| Direction |  | Circ. | Circ. | Circ. | Circ. | Circ. | Circ. | Circ. | Circ. |
| Rate Comparison |  |  |  |  |  |  |  |  |  |
| Female | 10 - 20 | - | - | - | - | - | - | - | - |
|  | 10 - 30 | - | + | - | - | - | - | - | + |
|  | 10 - 40 | + | + | + | + | + | + | + | + |
|  | 10 - 50 | + | + | + | + | + | + | + | + |
|  | 20 - 30 | - | - | - | - | - | - | - | - |
|  | 20 - 40 | + | + | + | + | + | + | - | - |
|  | 20 - 50 | + | + | + | + | + | + | - | - |
|  | 30 - 40 | - | - | - | - | - | - | - | - |
|  | 30 - 50 | + | - | - | + | + | - | - | - |
|  | 40 - 50 | - | - | - | - | - | - | - | - |
| Male | 10 - 20 | - | - | - | - | - | - | - | - |
|  | 10 - 30 | + | + | + | + | + | + | + | - |
|  | 10 - 40 | + | + | + | + | + | + | + | - |
|  | 10 - 50 | + | + | + | + | + | + | + | - |
|  | 20 - 30 | - | - | - | - | - | - | - | - |
|  | 20 - 40 | - | - | - | - | - | - | - | - |
|  | 20 - 50 | - | - | - | - | - | - | - | - |
|  | 30 - 40 | - | - | - | - | - | - | - | - |
|  | 30 - 50 | - | - | - | - | - | - | - | - |
|  | 40 - 50 | - | - | - | - | - | - | - | - |

Differences were observed for load rate intervals of 20-40 and 20-50 in the Equi-Biaxial protocol for females. Additionally, females exhibited different stretches in the Ratio-Y protocol for load rate intervals 10-30, 10-40, and 10-50 at a stress level of 30 kPa, while males showed no differences. In the Equi-Biaxial and Ratio-X protocols across all three stress levels, both females and males displayed differences between the 10-40 and 10-50 load rate groups. Furthermore, in the Equi-Biaxial and Ratio-X protocols, males exhibited differences compared to females in the 10-30 load rate comparison. Overall, females demonstrated more differences compared to males.

## 4 Discussions

Our biaxial testing confirms that human FPAs exhibit pronounced rate-dependent viscoelastic behavior. In both longitudinal and circumferential directions, faster loadings led to reduced stretch (i.e. a stiffer response), with the effect especially marked circumferentially. This finding underscores the time-dependent nature of arterial mechanics, consistent with the general viscoelastic principle that tissue stiffness increases as loading rate rises^5,25,26^. Notably, we found no significant sex-rate interaction, indicating that male and female FPAs respond similarly to varying loading rates. Although male samples showed slightly higher absolute compliance on average, this trend was modest and not statistically significant.

Overall, the rate sensitivity observed, particularly the circumferential stiffening at higher rates, highlights an important anisotropic viscoelastic response in FPAs. Our study employed three distinct loading protocols: Equi-Biaxial, Ratio-X, and Ratio-Y, each designed to assess different aspects of arterial behavior under varying loads. The Equi-Biaxial protocol, which applies equal force in both directions, revealed significant rate-dependent changes primarily in the circumferential direction. The Ratio-X and Ratio-Y protocols further emphasized this differential sensitivity. In the Ratio-X protocol, where the longitudinal load is halved, and the circumferential load is maintained, significant rate-dependent differences were again more pronounced in the circumferential direction. Similarly, the Ratio-Y protocol, with halved circumferential load, demonstrated significant rate dependency in the longitudinal direction, albeit to a lesser extent than the circumferential direction in the other protocols.

The greater rate-dependent stiffening in the circumferential direction of FPAs can be plausibly explained by the arterial microstructure and mechanobiology. Femoropopliteal arteries are muscular arteries with abundant smooth muscle cells (SMCs) oriented circumferentially in the medial layer^20,27^. Under rapid loading, these SMCs and the circumferentially aligned collagen fibers likely bear load quickly, allowing less time for viscoelastic relaxation, thus yielding a stiffer response. Indeed, vascular SMCs are known to respond to mechanical stimuli by generating actomyosin contractile forces that reduce compliance^28,29^. When arterial SMCs sense increased matrix rigidity or sudden stretch, they enhance their contractile tension, effectively stiffening the tissue^30^. Similarly, arterial SMC activation under mechanical stress further decreases arterial distensibility^28^. Although our experiments used *ex vivo* tissues with SMCs in a passive state, the structural alignment of SMCs and their residual tonus could contribute to the observed anisotropy. Future research is needed to explore the molecular and structural basis of the observed mechanical behavior due to increase in loading rate. This may provide deeper insights into the mechanisms driving viscoelasticity and loading-rate dependency in FPAs.

Our findings align with previous studies demonstrating rate-dependent mechanics in vascular tissues. For example, Anssari-Benam et al. showed that porcine aortic and pulmonary heart valve leaflets stiffen markedly with increasing biaxial stretch rate^26^. They found that even without preconditioning, the stress-stretch curves of valve tissue becomes stiffer at faster deformation rates, underlining the necessity of accounting for viscoelasticity in soft tissue testing. In arterial tissue, Stemper et al. reported that higher loading rates caused significant increases in the stress required to reach subfailure and ultimate failure in the porcine aorta, while the corresponding strains at failure decreased^25^. This behavior of higher stress and lower strain at failure is a clear indication of strain-rate-induced stiffening and is consistent with our observations in FPAs under sub-failure loads. Gaur et al. likewise observed a rate-dependent increase in failure stress in human diaphragm tissue, suggesting that many collagen-rich soft tissues become stiffer but less extensible when loaded rapidly^31^. These studies support the concept that biological tissues exhibit significant mechanical changes under varying loading rates.

Contrasting findings have been reported in other studies. For instance, Delgadillo et al. observed that pig thoracic aortas actually softened with increasing deformation rate, with loading forces reduced by up to ~20% when the strain rate was raised from 10%/s to 200%/s^32^. This inverse rate dependence, opposite to typical viscoelastic stiffening, suggests a complex interaction between arterial composition and loading conditions. Zemánek et al. found that loading rate had a negligible effect on the mechanical properties of the tissues of porcine thoracic aortas^10^. These discrepancies highlight that rate-dependent behavior can vary with vessel type and testing protocol. For example, experimental methods, such as load-controlled (force-rate) vs. displacement-controlled (strain-rate) testing, preconditioning routines, and specimen geometry (opened planar vs. intact cylindrical), can influence the observed rate effects. Our use of load-controlled biaxial tests (ramp in force at 10-50 mN/s) is one approach. Other studies using true strain-rate control or oscillatory loading might capture different viscoelastic responses. Nonetheless, the prevailing view from the literature and our results is that ignoring rate-dependency can lead to inaccurate characterization of arterial mechanics, as evidenced by multiple reports of significant viscoelastic effects in diverse vascular tissues.

The results presented here need to be considered within the study limitations. First, the range of loading rates (10-50 mN/s) was relatively narrow and did not include very slow or very fast loading regimes. Arterial viscoelasticity is known to be strain-rate dependent over several orders of magnitude. Thus, tests at lower rates (approaching quasi-static conditions) or higher rates (mimicking impact or pulsatile frequencies) could reveal additional aspects of FPA behavior. Second, the loading protocols were applied sequentially from low to high rate on each specimen. This order could potentially influence the results, as some tissue softening or additional conditioning might occur during the initial slow tests, affecting subsequent fast tests. While we preconditioned samples and did not observe obvious tissue damage, a randomized or reversed order of rate application would strengthen the validation of true rate effects unconfounded by prior loading history. Third, our sample size of 14 arteries and donor age range (45-55 years) were limited. We focused on mid-life adult specimens to reduce age variability, but arteries from younger and older individuals could exhibit different viscoelastic profiles due to developmental or degenerative changes in SMC function and ECM composition. Including a broader age range and a larger sample size would help determine how universally the observed trends apply, and whether factors like age or risk factors modulate rate-dependent stiffness^21^. We also note that because our experiments used excised arteries in a passive state, they did not account for active smooth muscle tone or live cellular responses. *In vivo*, vascular smooth muscle activation could dynamically change wall stiffness and how this active behavior interacts with passive viscoelasticity during rapid loading needs further investigation. Additionally, we focused only on one aspect of viscoelasticity, i.e., rate dependent mechanical behavior. To assess the complete viscoelastic properties of tissues, more advanced testing protocols, such as relaxation and creep tests, and models need to be designed^33,34^. Future research should aim to develop these tests, expand the sample size, and include a broader age range to validate and extend these findings.

## 5 Conclusion

This study demonstrates the significant loading-rate dependency of human FPAs, with more pronounced effects in the circumferential direction. These findings underscore the importance of considering viscoelastic properties and loading rates in the computational models relaying on arterial mechanical properties.

## 6 Acknowledgments

We wish to acknowledge Live on Nebraska for their help and support in collecting blood vessels and thank donors and their families for making this study possible. We also thank the Tissue Analysis Core (TAC) at the Center for Cardiovascular Research in Biomechanics (CRiB) for their assistance in collecting the blood vessels.

## 8 Ethics Statement

The human tissues used in this study were obtained from deceased donors through an organ procurement organization (Live On Nebraska) with informed consent provided by the next of kin. The collection and use of these tissues adhered to all relevant guidelines and regulations, including the Declaration of Helsinki. According to the U.S. Department of Health and Human Services and institutional guidelines, research involving cadaveric tissues does not constitute research on human subjects and, therefore, does not require Institutional Review Board (IRB) approval. Ethical approval was waived due to the nature of the study.

